# Difference in context-dependency between orienting and defense-like responses induced by the superior colliculus

**DOI:** 10.1101/729772

**Authors:** Kaoru Isa, Thongchai Sooksawate, Kenta Kobayashi, Kazuto Kobayashi, Peter Redgrave, Tadashi Isa

## Abstract

Previous electrical stimulation and lesion experiments have suggested that the crossed descending output pathway from the deeper layers (SCd) of superior colliculus (SC) controls orienting responses, while the uncrossed pathway mediates defense-like behavior. Here we extended these investigations by using selective optogenetic activation of each pathway in mice with channelrhodopsin 2 expression by double viral vector techniques. Brief photo-stimulation of the crossed pathway evoked short latency contraversive orienting-like head turns, while extended stimulation induced contraversive circling responses. In contrast, stimulation of uncrossed pathway induced short-latency upward head movements followed by longer-latency defense-like behaviors including retreat and flight. The novel discovery was that the evoked defense-like responses varied depending on the environment, suggesting that uncrossed output can be influenced by top-down modification of the SC or its downstream. This further suggests that the SCd-defense system can be profoundly modulated by non-motor, affective and cognitive components, in addition to direct sensory inputs.

## Introduction

The midbrain superior colliculus (SC) is an evolutionary ancient brainstem center controlling the initial, rapid, sensory-driven appetitive or defensive movements that are essential for survival in the natural environment (Dean et al., 1989). Its superficial layers (SCs) receive glutamatergic visual inputs directly from the retina (Isa et al., 1998) or indirectly from the visual cortex. Efferent projections from the SCs are directed to the visual thalamus (Lane et al., 1974), nucleus parabigeminalis (Sherk, 1979; Edwards, 1980; Bennett-clarke et al., 1989), or to deeper layers of the SC (SCd) (Isa et al., 1998; Lee et al., 1997; Doubell et al., 2003). The SCd receives non-visual sensory inputs and also contextual, motor or attention-related signals from forebrain regions (Wurtz and Albano, 1980; Sparks, 1986). The SCd neurons integrate information from these multiple sources and send efferent motor commands to the brainstem and spinal cord (Huerta and Harting, 1982), biologically salient visual cue signals to the midbrain dopamine neurons (Comoli et al., 2012), fear-related signals to the periaqueductal gray (Evans et al., 2018), and ascending motor efference copy signals to the thalamus (Sommer and Wurtz, 2006). In terms of behavioral control, previous studies in rodents suggested there are two major output channels that descend from the SC to the brainstem (Dean et al., 1989; Sahibzada et al., 1986). One crosses the midline in the ventral tegmental decussation and projects caudally via the predorsal bundle to the contralateral medial ponto-medullary reticular formation and upper cervical spinal cord. In rodents, this pathway mainly originates from the lateral part of the SC that represents the lower half of the contralateral visual field, and controls orienting and approach behavior (Dean et al., 1986). In carnivores and primates, saccadic eye movements and orienting head movements are controlled by this system (Wurtz and Albano, 1980; Sparks, 1986, 1999; Isa and Sasaki, 2002). The other descending projection, the ipsilateral one, originates mainly from neurons in the rostro-medial part of the rodent SC, which represents the upper visual field. It projects caudally to the cuneiform nucleus, and the ventrolateral ponto-medullary reticular formation (Redgrave et al., 1987, 1988). This pathway was considered to control defense reactions, including freezing and escape behaviors (Sahibzada et al., 1986). Previous studies investigating these SC’s descending pathways used electrical stimulation and lesion techniques combined with standard anatomical tract tracing and c-fos immunohistochemistry, to verify the location of activated neurons (Dean et al., 1989; Redgrave et al., 1987; King et al., 1996). However, functional analyses using these traditional, but cellularly nonselective techniques, ran potential risks of activating or lesioning passing fibers from the other projection, coupled with possible problems of post-lesion neuroplasticity. Furthermore, it has also been difficult to stimulate either the SCs or SCd neurons separately without affecting the other. The present study sought to overcome these limitations and investigate in more detail the cellular identity of orienting and defense systems in the SC and how they might be influenced by behavioral context. To this end, we tested the selective activation of the individual SC output pathways by combination of two viral vectors. One is the highly efficient retrograde gene transfer vector, a modified lenti-viral vector (Kato et al., 2011) carrying Cre, while the other is the adeno-associated viral (AAV) vector carrying channelrhodopsin 2 (ChR2) located downstream of a double flox sequence. This combination enabled a pathway-specific expression of ChR2. These procedures allowed us to activate each of the two SC-brainstem output channels independently. Furthermore, this double viral vector technique can reveal the morphology of the manipulated neurons, including cell bodies and axonal arborization, as shown in our previous studies using similar constructs (Kinoshita et al., 2012; Sooksawate et al., 2013; Ishida et al., 2016; Tohyama et al., 2017; Kinoshita et al., 2019). The present investigation has made extensive analysis of the relationship between the stimulus parameters and induced behavior in different environments, clarified status of the two major output pathways, and shown how they can be influenced by the environment.

## Results

### Pathway-selective optogenetic activation of superior colliculus (SC) output neurons

Procedures were developed to independently activate either the crossed or uncrossed descending pathways from the SC. For this purpose, one group of mice was injected with the retrograde gene transfer vector, NeuRet-MSCV-Cre into the left medial ponto-medullary reticular formation (PMRF) and AAV-DJ-EF1α-DIO-hChR2(E123T/T159C)-EYFP into the right (contralateral) SC (Figure 1A, C). In these mice, ChR2 was expressed exclusively in SC neurons projecting to the crossed descending pathway. In the second group of mice, the former vector was injected into the right cuneiform nucleus (CnF) and the second vector was injected into the right (ipsilateral) SC (Figure 1C). In these animals, the expression of ChR2 was restricted to the neurons whose axons projected to the uncrossed descending pathway. After the behavioral testing, electrophysiological experiments were conducted (Figure 1B). Electrophysiological recording in both groups showed that when optical stimulation with blue laser (473 nm wavelength) was applied to the right SC, either the crossed or the uncrossed output neurons in the SC were robustly activated (Figure 1D). The latencies of the spiking responses after the laser onset were 8.01 ± 0.66 ms (n = 23) for the crossed pathway neurons and 9.27 ± 1.54 ms (n = 13) for the uncrossed pathway neurons. Following repetitive optical stimulation (10 - 20 trains of 50 ms pulses at 10 Hz) both types of SC neurons were reliably activated.

**Figure 1.**
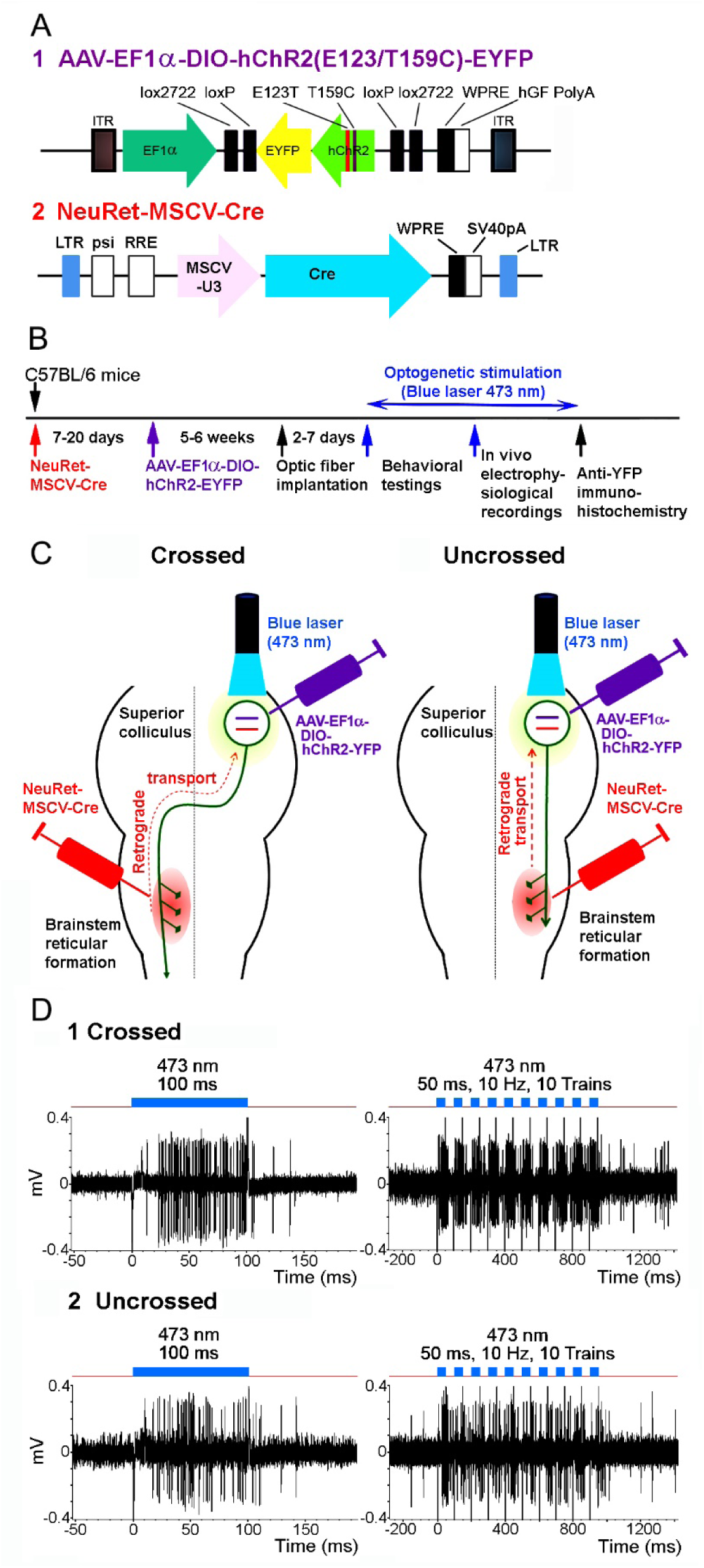
Pathway-selective optogenetic activation of SC output neurons. **(A)** Viral vector constructions. **(B)** Experimental protocols. **(C)** Diagram for the double injection of the viral vectors into the brainstem and SC, and the interaction of NeuRet-MSCV-Cre and AAV-EF1α-DIO-hChR2(E123T/T159C)-EYFP in the double infected SC neurons. **(D)** Responses of a mouse SC output neuron to blue laser stimulation with a 500 µm diameter fiber. (**1**) A neuron consisting the crossed pathway. (**2**) A neuron consisting the uncrossed pathway.

For the systematic evaluation of behavioral responses evoked by optical stimulation, they were classified as either contraversive orienting-like responses (horizontal head turn or circling), or defense-like responses (upward head turn, retreat, flight or freezing) (Figure 2A-D). The orienting-like responses were subdivided further into four categories reflecting the magnitude of responses through which contralaterally-directed head movements spanned; category 1: >180° circling, category 2: 90°-180° circling, category 3: <90° circling, category 4: head-only movement (Figure 2A-B). On the other hand, defense-like responses were more complicated. They often began with a quick upwardly directed head-only turn (category 4), and was often followed by backward walking (termed “retreat”, category 5, Figure 2C) and/or fast forward running away (termed “flight”, category 6, Figure 2D). Thus, smaller numbers below 4 indicated larger orienting, while the larger numbers above 4 were assigned for stronger defense-like responses.

**Figure 2.**
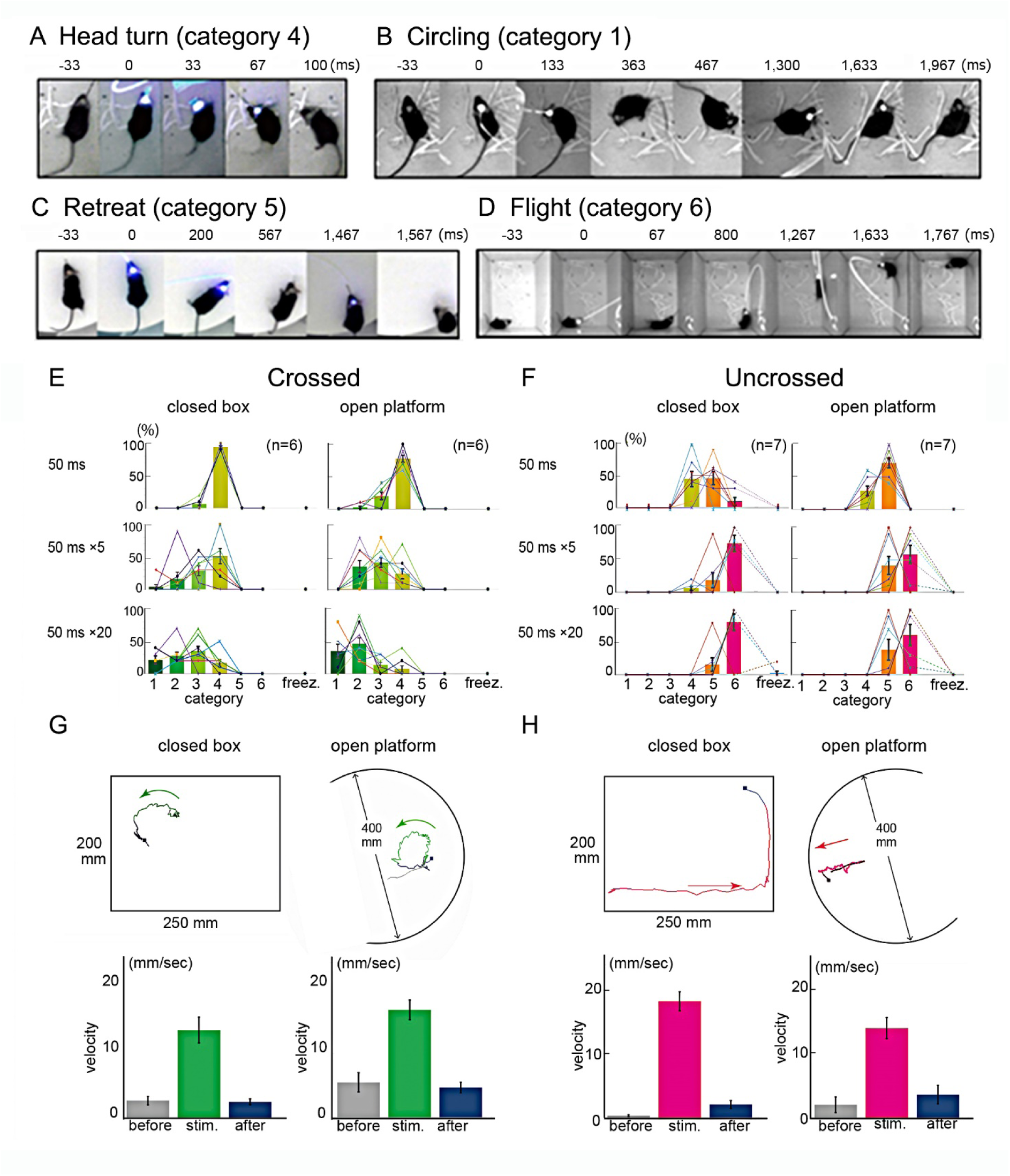
Categorization of behavioral responses to selective optogenetic activation of each SC output pathway. (**A-D**) Sequential photographs illustrating each of typical behavioral response patterns of the mice following optogenetic activation of SC neurons **(E)** Crossed SC-brainstem pathway. **(F)** Uncrossed SC-brainstem pathway. **(G)** Upper panel shows the examples of the trajectory of the defense-like responses of the mouse during stimulation of the crossed SC-brainstem pathway (20 trains of 50 ms pulses at 10 Hz) in the closed box and on the open platform. Lower panel shows a histogram of the velocity of body movements before, during and after stimulation. **(H)** Upper panel shows the examples of the trajectory of the defense-like responses of the mouse during stimulation of the uncrossed SC-brainstem pathway (20 trains of 50 ms pulses at 10 Hz) in the closed box and the open platform. Lower panel shows a histogram of the velocity of body movements before, during and after stimulation.

Evoked freezing responses were separately classified. To investigate the extent to which behavioral responses were context-dependent, observations were made either in a small box (25 cm × 20 cm), to which animals had been habituated for 3 hours prior to testing, or on a large open platform (an elevated circle 40 cm in diameter). Presentation of the different patterns of optical stimulation in the different testing environments were randomized.

Behavioral observations (Figure 2E) showed that when the crossed SC-brainstem pathway neurons were stimulated with a large optic fiber (500 µm in diameter) centrally located in the SC, brief stimulation (single pulses 50 ms) induced contraversive horizontal head turns in 94% of the trials in the closed box (category 4; Supplementary movie S1, n = 6). Progressively extended stimulation (200 ms pulses, or 5 - 20 trains of 50 ms pulses at 10 Hz) increased the frequency of circling with increasing amplitude (from category 3 to 2, and then to 1) (Figure 2E, 2G and Supplementary movie S2). When the stimulus duration was increased from single 50 ms pulses to 20 trains of 50 ms pulses at 10 Hz, average classification score decreased from 3.9 ± 0.03 to 2.5 ± 0.18 (p < 0.0001, Student’s t test). This confirmed that longer trains of optical stimulation evoked larger circling movements. All optically evoked movements were elicited at short latency (< 30 ms), almost always appearing in the first video frame following the laser onset. When the same stimulus was applied on the open platform, single 50 ms pulses induced head turn responses in 78 % of the trials, and when the stimulus duration was increased from single 50 ms pulses to 20 trains at 10 Hz, average classification score decreased from 3.8 ± 0.01 to 2.0 ± 0.02 (p < 0.0001, Student’s t test) (Figure 2E, 2G and Supplementary movie S3). No significant difference between the movement scores in the two environments could be found for both the brief stimulation (p = 0.47, Student’s t test) or the extended stimulation (p = 0.13, Student’s t test). Thus, similar contraversive responses were evoked in both testing environments. Consequently, these movements were considered to be context-independent.

When neurons projecting into the uncrossed SC-brainstem pathway were stimulated with brief optical stimulation (single 50 ms pulses) in the closed box, slightly ipsiversive upward head-only turns (category 4) and head turn plus retreat (category 5) were induced in 44% and 46 % of the trials, respectively (n = 7) (Figure 2F). Typically, the retreat responses were preceded by a small upward head turn, which appeared in the first video frame. With single pulse 200 ms stimulation, 73% of the responses were category 6 (flight), and category 5 (retreat) responses were induced in 20 % of the trials (Figure 2F and 2H, right). With extended stimulus duration (5 - 20 trains of 50 ms pulses at 10 Hz), the frequency of head-only turn responses became zero, while frequency of retreat (category 5) and flight responses (category 6) further increased (Figure. 2F and 2H, right) (average classification score increased from 4.7 ± 0.16 (single 50 ms pulses) to 5.7 ± 0.26 (20 trains of 50 ms pulses) (p = 0.012, Student’s t test)). The flight responses were directed mainly forward or sideways, not purely backwards, and looked like hiding the body in the corner. There was no fixed trend in direction of initial flight responses. An important novel finding was that the defense-like responses elicited by activation of the uncrossed SC-brainstem pathways were markedly context-dependent. In contrast, on the open platform, with the single 50 ms pulse, oblique-upward head-only turns (category 4) and head turn plus retreat (category 5) were induced in 29% and 71 % of the trials, respectively (Figure 2F). Here, the frequency of retreat responses was 71% and higher than in the closed box (p <0.05, Student’s t test, n = 7). With single pulse 200 ms stimulation, category 6 flight responses appeared in 23% of the trials, while category 5 retreat responses were induced in 69% of the trials (Figure 2E and 2G, right) (n = 7). With extended stimulus duration (5 - 20 trains of 50 ms pulses at 10 Hz), the frequency of head-only turn responses became zero, and frequency of retreat (category 5) also decreased to 39 % and the frequency of flight responses (category 6) increased to 61 %, where average classification score increased from 4.7 ± 0.07 (single 50 ms pulses) to 5.6 ± 0.16 (20 trains of 50 ms pulses) (p < 0.01, Student’s t test, n = 7). Thus, selective activation of the uncrossed SC-brainstem pathway was associated with induction of head-only turn (category 4), retreat (category 5) or flight (category 6) responses with infrequent freezing responses (in case of 20 trains of 50 ms pulses, freezing response was observed in only 5% in the closed box and 0% in the open platform, n = 7). In summary, in the closed box, stimulation with the same optical stimulation parameter frequently induced flight responses mainly consisting of moving forward from corner to corner (Figure 2G left and Supplementary movie S4), but on the open platform continuous retreat was the primary response (Figure 2G right and Supplementary movie S5). These data indicate that selective activation of uncrossed output from the SC was more associated with active defense-like movements at the expense of passive freezing responses. The induced movements were significantly affected by the environment in which the animals were placed.

### Axonal trajectories of neurons consisting the crossed and uncrossed SC-brainstem pathways

After the mice were sacrificed, the morphology of the activated cells was visualized with anti-GFP immunohistochemistry with diaminobenzidine (Figure 3). The somata of neurons consisting the crossed SC-brainstem pathway were concentrated in the most lateral portion of the SCd (Figure 3A3 and 3A9). A majority of the axons of these neurons crossed the midline in the ventral tegmental decussation and turned caudally to join the contralateral predorsal bundle (Figure 3A3-4). The axons continued into the medial ponto-medullary reticular formation (PMRF), where they approached the injection sites of the retrograde vector (Figure 3A5). Some fibers crossed the midline in the tectal commissure and terminated in the most lateral portion of the contralateral SCd, the mirrored location of the cells of origin (Figure 3A3, coSCd). The descending axons of these crossed neurons could be traced to pedunculo-pontine nucleus (PPN), and after crossing the midline, to the nucleus reticularis tegmenti pontis (NRTP), PMRF, raphe nuclei, inferior olive (IO) and spinal cord (Sp.c.) on the contralateral side (Figure 3A5-7). Some labeled axons could be traced to the SNc and in the ventral part of the ventral aspect of ipsilateral PMRF extending toward IO on the ipsilarteral side (Figure 3A4-6). In addition, the ascending collaterals could be traced to mediodorsal thalamic nucleus lateral part (MDL), lateral mesencephalic reticular formation (mRt), parafascicular thalamic nucleus (PF), ventral mediothalamic nucleus (VM), substantia nigra pars compacta (SNc), zona incerta (ZI) and rostral intralaminar thalamic nuclei such as paracentral thalamic nucleus (PC) and centrolateral thalamic nucleus (CL) on the ipsilateral side and central medial thalamic nucleus (CM) on both sides (Figure 3A1-2). The target areas of the crossed SC output cells are summarized in Figure 3A8.

**Figure 3.**
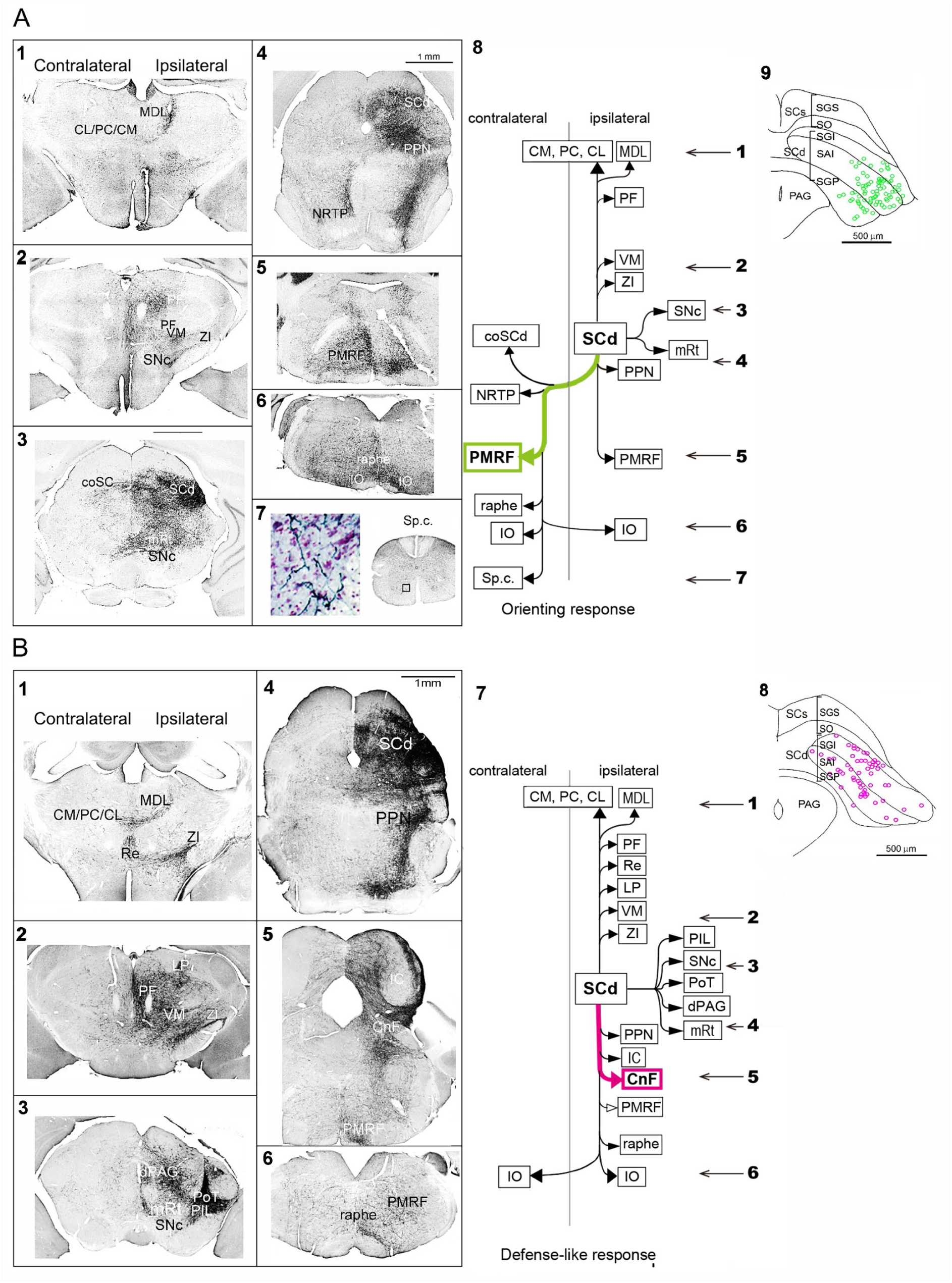
YFP-positive neurons and their axonal trajectories in the mice with double vector infection processed with anti-GFP immunohistochemistry. **(A)** Crossed SC-brainstem pathway; **1-7**. photomicrographs of the frontal sections of the diencephalon, brainstem and spinal cord, **8**. Targets of the crossed SC-brainstem neurons, with numerals with arrows indicating the rostrocaudal levels of photomicrographs in **1-7, 9**. A diagram showing the location of the somata of crossed SC-brainstem neurons. **(B)** Uncrossed SC-brainstem pathway; **1-6**. photomicrographs of the frontal sections of the diencephalon and brainstem, **7**. Targets of the uncrossed SC-brainstem neurons with numerals with arrows indicating the rostrocaudal levels of photomicrographs in **1-6, 8**. A diagram showing location of the somata of SC-brainstem neurons. **Abbreviations:** CM, PC, LC: rostral intralaminar thalamic nuclei such as central medial thalamic nucleus (CM), paracentral thalamic nucleus (PC) and centrolateral thalamic nucleus (CL), CnF: cuneiform nucleus, coSCd; contralateral SC deeper layers, dPAG: dorsal periaqueductal gray matter, IC: inferior colliculus, IO: inferior olive, LP: lateral posterior thalamic nucleus, MDL: mediodorsal thalamic nucleus lateral part, mRt: mesencephalic reticular formation, NRTP: nucleus reticularis tegmenti pontis, PF: parafascicular thalamic nucleus, PIL: posterior intralaminar nucleus, PMRF: medial ponto-medullary reticular formation, PoT: posterior thalamic nucleus, triangular, PPN: pedunculopontine nucleus, Re: reuniens thalamic nucleus, SAI: intermediate white layer, SC: superior colliculus, SCd: SC deeper layers, SGP: deep gray layer, SGS: superficial gray layer, SNc: substantia nigra pars compacta, SO: optic layer, Sp.c.: spinal cord, VM: ventral mediothalamic nucleus, ZI: zona incerta.

In contrast, although widely located throughout the rostral-caudal axis of the SCd, the somata of the neurons consisting the uncrossed SC-brainstem pathway were concentrated more medially (Figure 3B8) to those of the crossed SC-brainstem neurons. Consequently, there was little overlap between the location of the cell bodies of the crossed and uncrossed SC-brainstem pathways (c.f. Figures 3A9 and 3B8). The ascending axons of the uncrossed neurons could be traced to the CM, PC, LC, MDL, PF, reuniens thalamic nucleus (Re), lateral posterior thalamic nucleus (LP), VM, ZI and posterior intralaminar thalamic nucleus (PIL) (Figure 3B1-2).

At the same level of the SC, the axons projected to the SNc, posterior thalamic nucleus triangular (PoT), dorsal periaqueductal gray matter (dPAG) and lateral mesencephalic reticular formation (mRt) (Figure 3B3-4). The descending axons were projected to the inferior colliculus (IC), CnF, PPN and PMRF on the ipsilateral side and bilaterally to the IO (Figure 3B5-7). The target areas of the uncrossed SC output cells are summarized in Figure 3B7.

These results not only confirmed the previously described connectivity patterns of the crossed and uncrossed descending projection of the SC (Redgrave et al., 1987), but also found some novel projections not described in the previous studies, such as ipsilaterally descending branches of crossed SC-brainstem neurons targeting ipsilateral SNc, PPN, ventral part of PMRF, and re-crossed projection to IO (see Figure 3A8).

### Electromyogram (EMG) responses of dorsal neck muscles following optical stimulation of the crossed and uncrossed SC-brainstem pathways

To further investigate the synaptic connectivity between the crossed and uncrossed SC-brainstem pathways and spinal motoneurons, we recorded the EMG responses of dorsal neck muscles evoked by ipsilateral or contralateral stimulation of the SC in anaesthetized animals (Figure 4A). When the crossed SC-brainstem pathway was activated, the mean latency of evoked EMG responses in contralateral (left) muscles was 12.85 ± 0.96 ms (n = 6), and in ipsilateral (right) muscles was 13.01 ± 1.17 ms (n = 6) (Figure 4B). Considering the previous observation that spiking activity in crossed SC output neurons was induced at the latency of 8.01 ± 0.66 ms (n = 23) after the laser onset (Figure 4D1), the conduction time from the SC to the neck muscles must have been in the range of 4-5 ms, on both sides. These ultra-short latency muscular responses were completely abolished after unilateral microinjections of 0.1 μl muscimol (0.1 – 1.0 mg/ml) into the left PMRF (Figure 4B), the injection sites of retrograde vector. This result suggests that activation of the dorsal neck muscles by the SC is mediated primarily by tecto-reticulo-spinal pathways with relay neurons in the PMRF which have uncrossed or crossed connections to neck motoneurons (Isa and Sasaki, 2002). Following activation of the uncrossed SC-brainstem pathway (Figure 4C), the mean latency of EMG responses in the contralateral (left) muscles was 13.52 ± 1.04 ms (n = 7), and in the ipsilateral (right) muscles was 12.41 ± 1.13 ms (n = 7). Considering again that the average latencies of optically evoked spiking responses of the relevant SCd neurons was 9.27 ± 1.54 ms (n = 13) (Figure 4D2), the conduction time in the uncrossed descending projection from the SC to the neck muscles was about 3-4 ms. Again, these short latency EMG responses were abolished after microinjections of muscimol into the brainstem close to the right CnF (Figure 4C), the injection site of retrograde vector. These results suggest that the excitatory drive from the SC to the dorsal neck muscles mediated via the uncrossed SC-brainstem pathway have a critical relay in the brainstem area close to the CnF, presumably reticulospinal neurons in the PMRF with crossed or uncrossed descending projections to the neck motoneurons (Isa and Sasaki 2002). Taken together, the short latencies of the evoked EMG responses suggest that the synaptic connectivity from the SC to neck muscles via both crossed and uncrossed pathways are minimally disynaptic or oligosynaptic and mediated by the reticulospinal neurons in PMRF.

**Figure 4.**
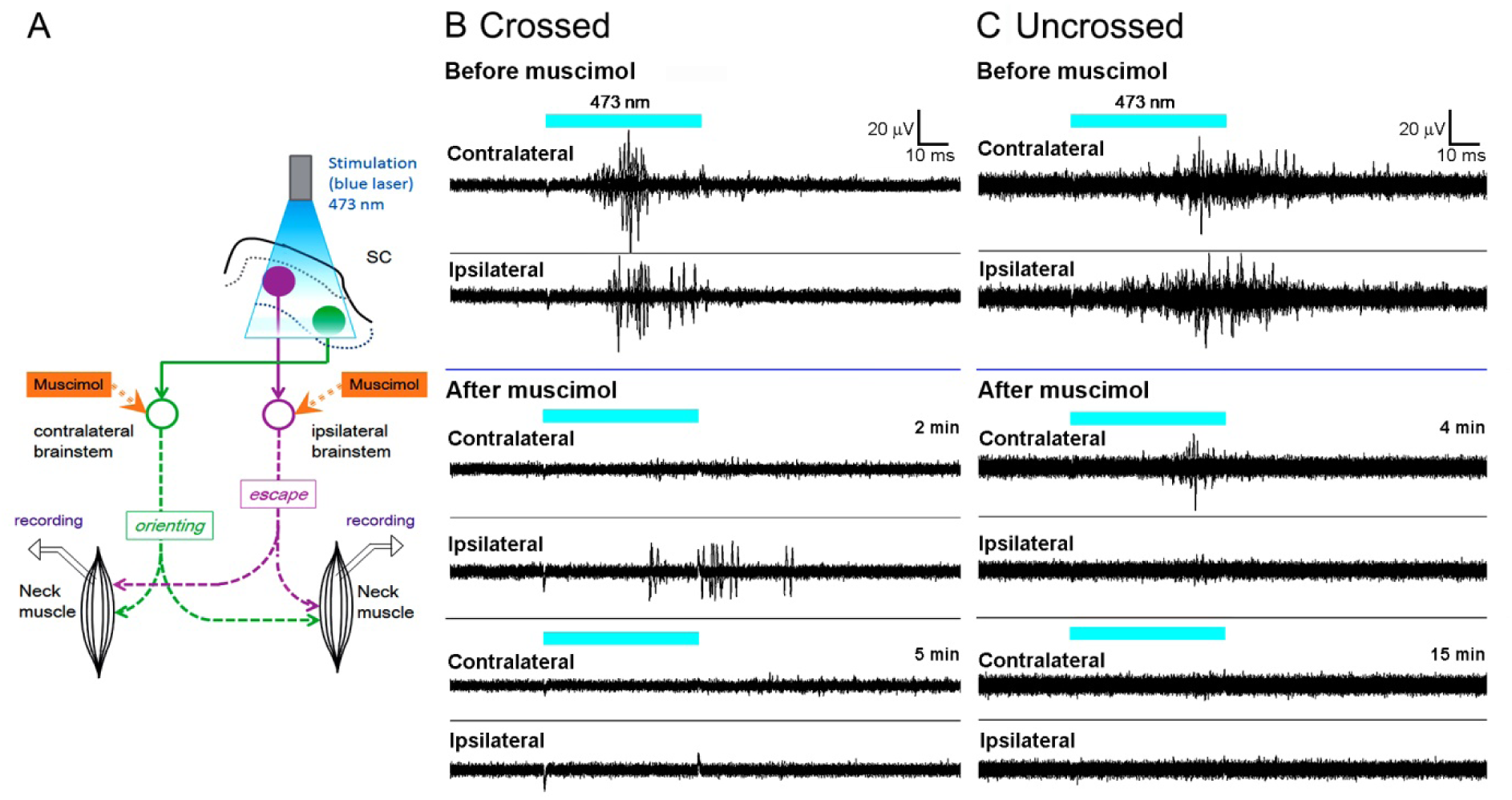
EMG responses of dorsal neck muscles to optogenetic stimulation of the SC neurons. **(A)** Diagram showing the experimental protocol. **(B)** EMG response to the activation of the crossed SC-brainstem pathway, which was inhibited by a muscimol injection into the brainstem. **(C)** EMG responses to the activation of the uncrossed SC-brainstem pathway, which were also abolished by unilateral microinjection of muscimol into the injection site of the NeuRet vector in the brainstem.

### Nonselective optogenetic activation of the SC neurons

To compare the effects of pathway-selective activation with those of nonselective activation of the SC, AAV-CAG-Ch2R(H134R)-tdTomato was injected into the SC. As shown in Figure 5A, neurons in virtually the whole SC, including both SCs and SCd, expressed ChR2-tdTomato. Optical stimulation with blue laser was applied at 3 locations (Figure 5B) in SC; (1) a 250 μm diameter fiber was used to stimulate neurons representing the lower visual field in the caudo-lateral SC, (2) the same 250 μm diameter fiber was used to activate the rostro-medial aspect of the SC representing the upper visual field, and (3) a larger 500 μm diameter fiber was placed in the central part to activate the wide area of the SC. Under anesthesia, the neurons in the SCd located below the optic fiber were reliably activated by the laser stimulation, in response to both prolonged pulses of laser stimulation (100 ms duration) and repetitive trains of short pulses (10 trains of 50 ms pulse at 10 Hz) (Figure 5C). Brief optical stimulation (single 50 ms pulses) of the caudo-lateral SC induced contraversive orienting responses of the head (category 4) in the awake condition (Figure 6A). With the extended stimulation, large angular rotations of the head eventually followed by contraversive circling were observed (from category 3 to 2, and then 1). When the stimulus duration was increased from single 50 ms pulses to 20 trains of 50 ms pulses at 10 Hz (Figure 6A), the average of the classification score reliably and systematically decreased from 4.0 ± 0.03 (mean ± SE) to 2.1 ± 0.25 (p < 0.0001, Student’s t test) in the closed box and from 4.0 ± 0.03 to 2.1 ± 0.26 (p < 0.0001, Student’s t test) on the open platform. No significant difference between the scores of the two environments could be found for both the brief stimulation (p = 0.49, Student’s t test) and the extended stimulation (p = 0.47, Student’s t test). Thus, the evoked responses were context-independent.

**Figure 5.**
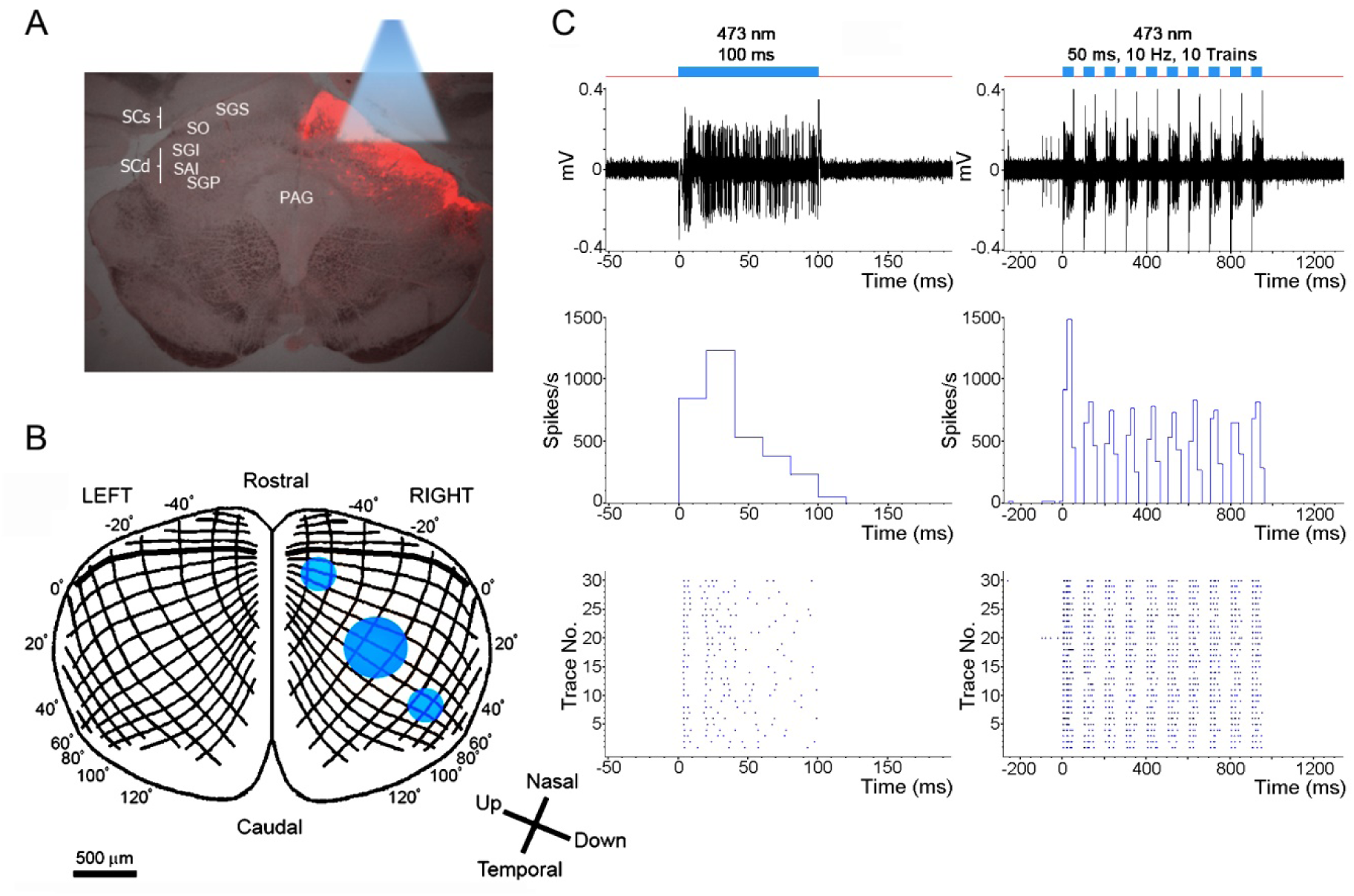
Nonselective optogenetic activation of the SC neurons. **(A)** Fluorescent photomicrograph of the superior colliculus that expressed tdTomato. **(B)** Site of laser stimulation on the topographic map of the mouse SC. **(C)** Responses of mouse SCd neuron to blue laser stimulation; raw traces (upper row), peristimulus time histogram (PSTH) (middle row) and raster plots (lower row). Single, long pulse (100 ms) on the left, and repetitive stimulation with short pulses on the right (10 trains of 50 ms duration pulses with 50 ms interval (10 Hz)). **Abbreviations:** PAG: periaqueductal gray matter, SAI: intermediate white layer, SC: superior colliculus, SCd: SC deeper layers, SCs: SC superficial layers, SGI: intermediate gray layer, SGP: deep gray layer, SGS: superficial gray layer, SO: optic layer.

**Figure 6.**
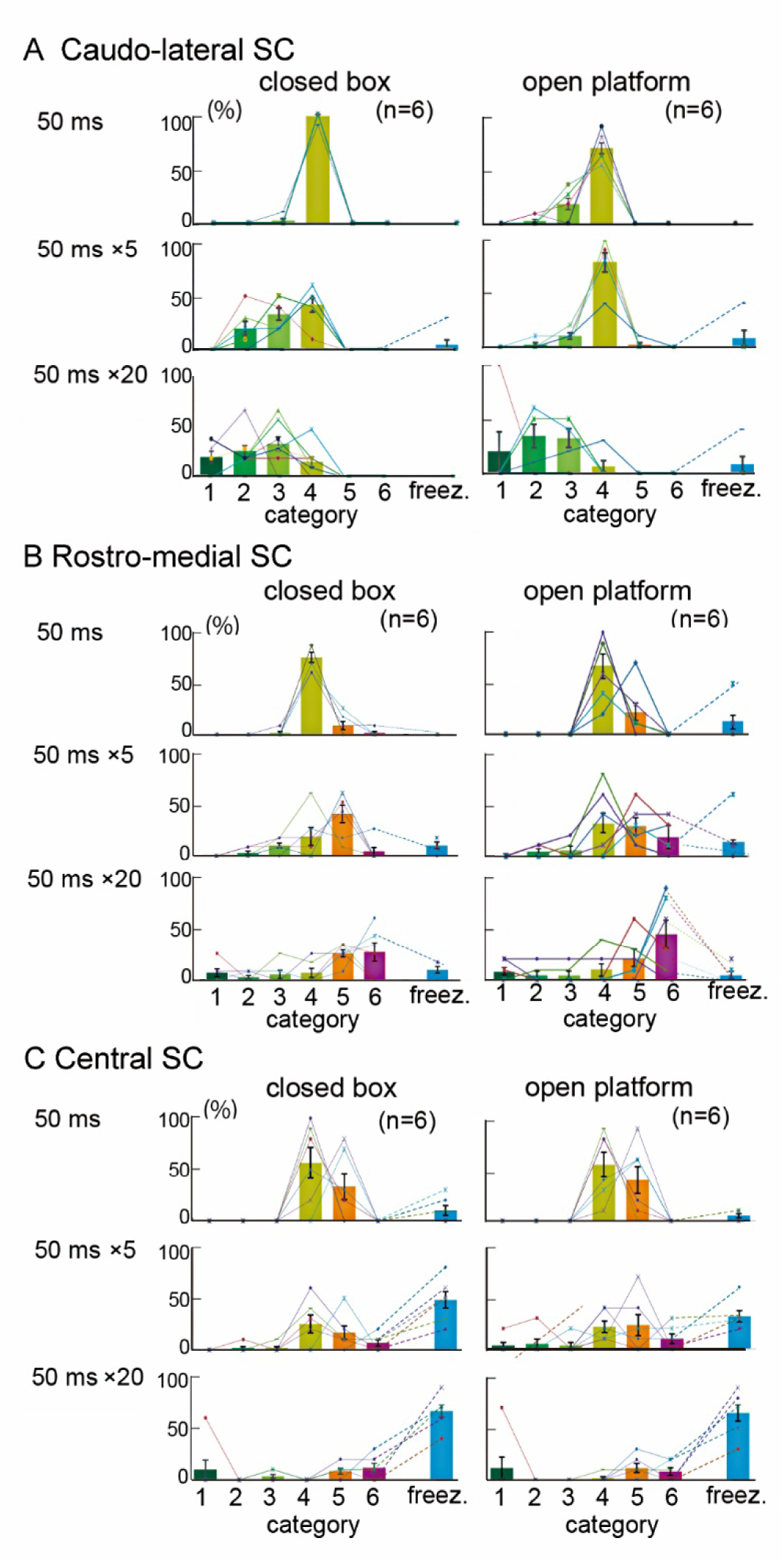
Categorization of behavioral responses to nonselective optogenetic activation of the SC neurons in the closed box (left) or in open platform environment (right). **(A)** Stimulation of the caudo-lateral SC with a 250 µm diameter fiber. **(B)** Stimulation of the rostro-medial SC with a 250 µm diameter fiber. **(C)** Stimulation of the central SC with a 500 µm diameter fiber. Behavioral response categories (category 1: >180°, category 2: 90°-180°, category 3: <90°, category 4: contralateral head-only turn responses, category 5: retreat, and category 6: flight, and freezing responses) are plotted on the horizontal axis and their percentage of occurrence to stimulation with each parameter on the vertical axis.

In case rostro-medial SC stimulation was applied in the closed box, brief stimulation (single 50 ms pulses) tended to induce mainly an upward and slightly ipsiversive head turn (on average, 87%, category 4, n = 6). When the pulse width was increased (5 trains of 50ms pulses at 10 Hz) the frequency of retreat responses increased (47%, category 5). If the stimulus intensity was increased further (20 trains of 50 ms pulses at 10 Hz), the stimulation frequently evoked flight responses (32%, category 6). On the open platform, the extended stimulation (5-20 trains of 50 ms pulses at 10 Hz) often induced flight responses (45%, category 6). Here, the responses to nonselective activation of the rostro-medial SC and pathway-selective activation of uncrossed SC-brainstem pathway (Fig. 2F) were somewhat different, most likely because of the difference in the diameter of optic fibers (250 μm diameter in the former cases and 500 μm in the latter cases) or because of pathway-specificity of the stimulation.

When the wide area of the SC was stimulated by the 500 μm diameter optic fiber placed over the central SC, a short stimulation (50ms) evoked head turn (55% in closed box and 57% in open platform), (Figure 6C). With the increases in the duration of optical stimulation (20 trains of 50ms pulses at 10 Hz), the frequency of freezing response markedly increased (on average, 67% in the closed box and 65% on the open platform, n = 6) and retreat (category 5) and flight (category 6) responses were induced less frequently (on average, categories 5 and 6 responses were 8% and 12% in the closed box, respectively, and those were 12% and 8% on the open platform, n = 6) (Figure 6C). We classified the responses mainly by the major motor responses, however, in many cases, these responses were typically followed by freezing in the case of central SC stimulation.

Thus, defense-like responses were evoked most frequently from local stimulation of the rostro-medial part of the SC, representing the upper visual field, while local stimulation of the caudo-lateral SC, representing the lower visual field typically elicited orienting responses. Furthermore, freezing was the most notable feature of the non-specific stimulation of the wide area of SC.

## Discussion

In this study, we expressed ChR2 selectively in each of the two major descending output channels from SCd and the behavioral effects of activating each output cell group were investigated. Pathway associated selectivity was achieved by combining two different viral vectors, one of which was injected into the location of the cells of origin in the SC and the other into their respective target areas in the brainstem. This technique can be applied to any anatomically identified pathway, even if we do not know the cell-type specific promotor. Brief optical activation of crossed SC-brainstem pathway caused contraversive orienting-like head turn, while extended stimulation caused contraversive circling involving the whole body. In clear contrast, optical activation of the uncrossed SC-brainstem pathway elicited defense-like responses. In these cases, brief stimulation induced oblique and slightly ipsiversive head-only turns directed to the upper visual field. However, with extended stimulus duration, the short latency head turns were followed by responses resembling retreat or flight. These patterns of defensive responses were reminiscent of those described by Dean and Redgrave and colleagues in 1980’s, who studied the functional status of descending projections from the SC with classical electrical and chemical stimulation techniques combined with surgical lesions (Dean et al., 1989; Sahibzada et al., 1986; Dean et al., 1986; Redgrave et al., 1987, 1988; King et al., 1996).

We compared the effects of activating the descending projections selectively with those obtained following nonselective optical activation of SC neurons. Stimulation of the caudo-lateral SC, in which the lower visual field is represented, evoked responses that were similar to selective activation of the crossed SC-brainstem pathway. Alternatively, nonselective activation of the rostro-medial SC, where the upper visual field is represented, induced responses almost similar to those produced by selective activation of the uncrossed pathway. These results can be explained by considering the anatomical distribution of the cells of origin of the two descending pathways in the rodent SC, in conjunction with the ecological niche occupied by rats and mice (Dean et al., 1989). First, the neurons giving rise to the crossed descending projection are concentrated in the lateral SC in the mouse, while those projecting to the uncrossed pathway are more prevalent medially (Figure 3A9 and 3B8). In the retinotopic representation of visual space in the SC, medial region corresponds to the upper visual field, while the lower visual field is represented laterally. For these species, their predators (birds of prey and larger mammals) most frequently approach from above. It is therefore perhaps not surprising that in these species, a specialized association between upper field visual stimuli and defensive behavior has evolved. Alternatively, most stimuli that these animals would want to locate and move towards (food and young) are typically on the ground and would appear in the lower visual field. Clearly, the evolved mechanism for this functional specialization in rodents is the differential concentration of defense-related cells in the medial upper-field portion of the SC, and cells promoting orienting behavior located laterally in lower-field regions of the SC map (Redgrave et al., 1986).

A closer look at the results of the present study, however, reveals a more nuanced picture. Functional differences were observed between the nonselective activation of SC, which most likely involved the simultaneous activation of both descending projections and activation of the superficial layers (SCs). When the central SC was activated with an optic fiber with large diameter, freezing responses were observed more frequently than the localized stimulations. Optical stimulation with a large fiber could conceivably correspond to the sudden appearance of a large object covering much of the animal’s visual field, similar to the looming stimulus. Such a stimulus event would be unusual in nature, and it would be likely that both orienting-approach and defense-avoidance systems in the SC would have been activated simultaneously. If so, unresolved competition between them could mean that freezing would be the most adaptive response. An alternative possibility for the prevalence of the freezing being evoked by central SC stimulation could be that the non-pathway-selective optical stimulation may also have activated neurons in the SCs. Recent studies by Wei et al. (2015), and Shang et al. (2015), reported defense-like freezing following selective activation of cell groups in the SCs. This response was ascribed to indirect activation of amygdala by visual inputs routed through the SC (see below).

An important advantage of the current pathway-selective activation procedures was that the entire network of axonal projections, including collaterals, of activated neurons could be traced. Consequently, we were able to associate specific sets of behavioral reactions to identified anatomical substrates. In case of crossed SC-brainstem pathway neurons possess ascending branches to the midbrain and thalamic nuclei in addition to the caudal projection to the brainstem motor centers (Figure 3A8). Uncrossed pathways also possess ascending branches projected widely to a variety of midbrain and thalamic nuclei, some of which are different from the targets of the crossed pathway in addition to the descending branches (Figure 3B7). It is likely that the ipsilateral target regions may be associated with different components of the elicited defense-like responses. For example, the SC-PPN/CnF projections might be involved in the locomotor aspects of defense-like behaviors, because PPN and CnF have been considered as the mesencephalic locomotor region (Shik et al. 1966).

Recently, Wei et al. (2015) reported that CaMKII-positive neurons in the SCs were activated by looming stimuli and involved in freezing elicited by overhead threat (Wei et al. 2015). They further showed that the projections from the SCs to the LP and then to the amygdala were critical for the freezing response. Although the authors argued that the critical CaMKII-positive neurons were located in the intermediate layer, their figure showed the labeled neurons mostly in the SCs. Specifically, the wide field vertical cell group in this layer were implicated (Gale and Murphy, 2014). Also using modern tracing technology, the SC oligosynaptic projection to the amygdala was confirmed recently by Shang et al. (Shang et al. 2015). They found that the glutamatergic parvalbumin positive neurons in the SCs project to the amygdala indirectly via the parabigeminal nucleus (PBG). These parvalbumin positive neurons in the SCs were shown to be critical for the defensive responses elicited by looming visual stimuli. A more recent study by Shang et al. (2018) showed that the SC orchestrates the contributions of these two ascending pathways for defensive behaviors. Furthermore, Shang et al. (2019) showed that a group of SC-ZI pathway neurons is involved in predatory hunting. These neurons might comprise a subgroup of the crossed SC-brainstem pathway neurons, because the movements induced by activating these neurons were more characteristic of approach rather than escape. Further studies will be needed to understand how the neurons activated in these studies operate to influence the downstream hindbrain substrates for approach and defense. Certainly, from an evolutionary perspective, it would make more sense to have a threatening visual event capable of influencing appropriate motor output in the shortest possible time. It is therefore significant that we were able to show that activation of either of the SC’s principal descending pathways evoked EMG responses in the dorsal neck muscles about 3 - 5 ms after optically induced spiking in the SCd. The descending routes of communication were confirmed when critical relays of the uncrossed and crossed pathways in the brainstem reticular formation were inactivated, and optically elicited EMG responses abolished. This result is consistent with previous studies in cats that show the shortest uncrossed pathway from the SC to neck motoneurons is mediated by the reticulospinal neurons in the PMRF (Iwamoto et al., 1990; Anderson et al., 1971). It is known that there is a direct tectospinal projection, however, from the number of axons and boutons present at spinal levels its direct influence is likely to be weak (Isa and Sasaki, 2002). Therefore, it is more likely that the major tectal control of the neck motoneurons is mediated by a di- or oligosynaptic pathway through tecto-reticulospinal neurons. In case of the uncrossed pathway, the short latency EMG responses could be explained by a tri-synaptic linkage from the SC intermediate layer to neck motoneurons as shown previously in the cat by Alstermark et al. (1992).

In addition to the robust short latency responses, we also observed longer latency defense-like responses that appeared to be more context-dependent. It is therefore likely that context-dependent influences over defense-like responses in our study were mediated by the ascending collateral projections from uncrossed SC-brainstem pathway. These target regions of the thalamus including LP, MDL, PF, VM, and the intralaminar nuclei, have connections to the cerebral cortex, limbic system (Doron and Ledoux, 2000) and basal ganglia (Beckstead 1984). Return connections (direct and indirect) (Edwards, 1980; Wurtz and Albano, 1980; Redgrave et al., 1988; May, 2006) from these forebrain regions to the SC and other brainstem targets could be the means whereby affective and cognitive influences might modify the motor component of defensive responses to visual threat. In this regard, it is particularly interesting that the ascending collaterals from the SCd may reach the amygdala, because if such efference copy signal of the innate defense response activates the amygdala, it would indicate the existence of retrospective activation system of fearful emotion following the motor responses, differently from the visual feedforward activation of the amygdala shown in the recent studies of Shang et al. (2015, 2018) and Wei et al. (2015). Finally, a recent study by Evans et al. (2018) suggested that the SC pathway to the dPAG sets the threshold for the defensive behaviors, which in turn could be related to the context-dependency of the defense-like behaviors observed in the present study.

In summary, it is now clear that there are multiple output channels from the SC, from both SCs and SCd, which mediate approach orienting and defense-related signals. The precise nature of these responses can be modified by contextual cues. The role played by each channel in different aspects of the adaptive responses to appetitive and threatening visual stimuli, and how they can be modified by top-down influences, should be clarified in future studies.

## Acknowledgements

We thank Dr. K. Svoboda for providing the gene AAV-CAG-hChR2-H134R-tdTomato and Dr. K. Deisseroth for providing the gene AAV-EF1α-DIO-hChR2(E123T/T159C)-EYFP. This study was supported by AMED under Grant Number JP18dm0207020, a Grant-in Aid for Scientific Research on Innovative Areas (“Adaptive Circuit Shift” and “Hyper-adaptability”) and Kiban (A) from the MEXT Japan, and CREST “Opto-bio” Project from JST to T.I. We thank Yukio Nishimura for technical advice on EMG recordings, Toshie Kuwahara for assistance in the research and Masatoshi Kasai for a program for laser stimulation.

## AUTHOR CONTRIBUTIONS

K.I and T.S. equally contributed to this study by conducting the experiments. K.I., T.S., P.R. and T.I. designed the research, and wrote the manuscript. Kazuto K. and Kenta K. constructed the viral vectors. All authors contributed to the final version of this manuscript.

## STAR ⋆METHODS

### KEY RESOURCE TABLE

| REAGENT OR RESOURCE | SOURCE | IDENTIFIER |
| --- | --- | --- |
| Antibodies |  |  |
| Rabbit anti-GFP antibody | ThermoFisher,<br>Japan | Cat# A11122 |
| Biotinylated goat anti-rabbit IgG | Vector<br>Laboratories, USA | Cat# BA1000 |
| ABC-Elite | Vector Laboratories, USA | Cat# PK6100 |
| Recombinant DNA |  |  |
| NeuRet-MSCV-Cre | Kazuto Kobayashi | N/A |
| pAAV-CAG-hChR2(H134R)-tdTomato | Karel Svoboda | N/A |
| pAAV-EF1α-DIO-hChR2(E123T/T159C)-EYFP | Karl Deisseroth | N/A |
| Software and Algorithms |  |  |
| EthoVision XT 11.5 | Noldus, Netherland | N/A |
| pClamp 10.2 | Molecular Devices, CA, USA | N/A |
| Others |  |  |
| Plastic optical fiber | Lucir Inc., Tsukuba, Japan | N/A |
| Laser source (LUCIR, Model COME2-OFC-1) | Lucir Inc., Tsukuba, Japan | N/A |
| Flash stimulator (SLS-3100) | Nihon Kohden, Japan | N/A |
| Microelectrode amplifier (MEZ-8301) | Nihon Kohden, Japan | N/A |
| Analog-to-digital converter (Digidata 1440A) | Molecular Devices, CA, USA | N/A |

### LEAD CONTACT AND MATERIALS AVAILABILITY

Further information and requests for resources and reagents should be directed to and will be fulfilled by the Lead Contact, Tadashi Isa

## Materials and methods

### Animals

Forty-two 10-week-old male C57BL/6 mice were used in this study. The experimental protocol for the use of animals was conducted following the Guidelines of the National Institutes of Health and the Ministry of Education, Culture, Sports, Science and Technology (MEXT) of Japan and approved by the Institutional Animal Care and Use Committee of the National Institutes of Natural Sciences. We made all attempts to minimize the stress, distress and number of animals used.

### Plasmid construction

The plasmid for NeuRet-MSCV-Cre vectors was obtained from a DNA fragment encoding the Cre recombinase gene (Kobayashi et al., 2004) and the murine stem cell virus promoter of pCL20c-MSCV (Hanawa et al., 2002; 2004). The pAAV-CAG-hChR2(H134R)-tdTomato and pAAV-EF1α-DIO-hChR2(E123T/T159C)-EYFP were provided from Drs. K. Svoboda and K. Deisseroth, respectively.

### Viral vector preparation

NeuRet vectors were prepared as described previously (Kato et al., 2014). The copy number of RNA genome was estimated by a Lenti-X qRT-PCR titration kit (Clontech, Palo Alto, CA). Real-time quantitative PCR was performed in duplicate samples using the StepOne real-time PCR system (Applied Biosystems, Tokyo, Japan). AAV vectors were packaged as described previously (Kobayashi et al., 2016). The copy number of viral genome (vg) was determined by the TaqMan Universal Master Mix II (Applied Biosystems, Foster City, CA).

### Viral vector injection

For the pathway-selective expression of ChR2, NeuRet-MSCV-Cre (0.4-0.5 µl; titer, 1.1-30 × 10^11^ copies/ml) and AAV-EF1α-DIO-hChR2(E123T/T159C)-EYFP(0.15-0.5 µl; titer, 4.5 × 10^10^ vg/µl) were injected into the target area of SC-brainstem neurons and the location of the pathway cells of origin, respectively. For the contralateral pathway group, injection of NeuRet-MSCV-Cre into the left medial pontine reticular formation inclined by 45 degrees caudally (Flanklin and Paxinos 2007) and that of AAV-EF1α-DIO-hChR2(E123T/T159C)-EYFP were also described before (Sooksawate et al., 2013). For the ipsilateral pathway group, NeuRet-MSCV-Cre was injected into the right brainstem close to the cuneiform nucleus inclined as above, −8.2-8.7 mm from Bregma, 1.2 mm lateral to the midline.

The AAV-EF1α-DIO-hChR2(E123T/T159C)-EYFP was injected into the right SC with the same coordinates.

Viral vector injection for nonselective optogenetic activation into the right SC was performed as described before (Sooksawate et al., 2013). AAV-CAG-hChR2(H134R)-tdTomato (1.9 × 10^9^ vg/µl) was injected as described above.

### Optic fiber implantation

Four to seven weeks after the viral injection(s), mice were anesthetized as mentioned above. Then, the 250-µm or 500-µm diameter plastic optical fiber (Lucir Inc., Tsukuba, Japan) was implanted through the cortex just above the SC surface using stereotaxic procedures. For the selective activation of output neurons of the SC, a 500-µm diameter optical fiber was implanted through the cortex just above the SC surface with the coordinate of −3.8 mm from Bregma, 1.2 mm right to the midline and 0.8-1.0 mm from the surface of the cerebral cortex. For the nonselective activation of the SC neurons, we divided the mice into 3 groups, caudo-lateral, rostro-medial, and central SC. 500-µm optical fiber was implanted in the “central SC” group and 250-µm optical fiber was implanted into “caudo-lateral” and “rostro-medial” groups. The stereotaxic coordinates for the caudo-lateral SC was −4.0 mm from Bregma, 1.6 mm right to the midline and 0.8-1.0 mm from the surface of the cerebral cortex, that of the rostro-medial SC was −3.4 mm from Bregma, 0.8 mm right to the midline and 0.8-1.0 mm from the surface of the cerebral cortex, and that of the central SC was the same coordinates as of the selective pathway activation.

### Behavioral effects with laser stimulation

Two to seven days after the mice recovered from optical fiber implantation, they were placed into a closed box (20 cm wide, 25 cm long and 30 cm high) or on an open elevated circular field (open platform, 40 cm diameter and 1 m high from the floor) for testing the effect of blue laser illumination from a laser source (LUCIR, Model COME2-OFC-1, Lucir Inc., Tsukuba, Japan). The intensities for laser stimulation were 20-170 mW/mm^2^ for 250-µm diameter optical fiber and 36-260 mW/mm^2^ for 500-µm diameter optical fiber. We stimulated the SC neurons with a single pulse of 50-200 ms duration or repetitive stimulation with 50 ms duration and 100 Hz frequency for 0.5-2.0 s. Behavioral data were analyzed with EthoVision XT 11.5 (Noldus, Netherland). Stimulation with individual parameters were repeated 10 times in each animal. Animals were randomly grouped. The numbers of mice used in each group were 6-7.

### *In vivo* electrophysiological recordings and EMG recordings

After the behavioral experiments, 3-4 mice of each group were anesthetized with urethane (1.2-1.5 g/kg i.p.). Dexamethasone (5.5 mg/kg) and atropine (0.1 mg/kg) were injected intramuscularly. Their heads were fixed to the stereotaxic apparatus. Optical fibers implanted for the testing and a small patch of the skull were removed to expose the cortex overlying the SC. The right eye was covered by plastic tape to protect against the light passing through this eye. 250-µm optical fiber was lowered vertically into the brain with the same coordinate as the removed optical fiber. A thin tungsten electrode (2 MΩ resistance) was inclined by 20 degrees and placed at 0.4 mm rostral to the rostral edge of optical fiber at the surface of the cortex and lowered into the right SC using a motor drive manipulator (DM System, Narishige, Japan). To confirm the location of the right SC, light flashes (Flash stimulator, SLS-3100, Nihon Kohden, Japan) was used to stimulate the left eye and monitor the light evoked potential in the right SC. The depth of SCd was identified by observation of the reversal of visually guided field potential (Katsuta and Isa, 2003), which was amplified by microelectrode amplifier (MEZ-8301, Nihon Kohden, Japan). Data were digitized by analog-to-digital converter (Digidata 1440A, Molecular Devices, CA, USA) and acquired using a pClamp system (pClamp 10.2; Molecular Devices, CA, USA).

For EMG recordings, after behavioral test, the mouse was anesthetized with a ketamine-xylazine combination (60 mg: 10 mg/kg i.p.). Dexamethasone (5.5 mg/kg) was injected intramuscularly. Then, the mouse head was fixed to the stereotaxic apparatus. The EMG electrodes, stainless needles, were inserted into the neck on both sides. The EMG responses were measured during blue laser illumination from these muscles before and after injection of GABAA receptor agonist, muscimol (0.1 µl, 0.1-1 mg/ml) into the injection sites of NeuRet-MSCV-Cre, left medial pontine reticular formation or right CnF using the same stereotaxic coordinates and method as the vector injections. The numbers of mice used in each group were 6-7.

### Immunohistochemical assessments

At the end of the experiments, the mice were deeply anesthetized with intraperitoneal injection of sodium pentobarbital (80 mg/kg body weight) and transcardially perfused with 0.05 M PBS followed by 4% paraformaldehyde in 0.1 M phosphate buffer (pH 7.4). The brain and spinal cord were cryoprotected and sectioned at a thickness of 40 μm using a sliding microtome (Retoratome REM-710, Yamato, Asaka, Japan). The expression of ChR2-YFP in the double infected SC output neurons, including their somata and axons, were examined with anti-GFP immunohistochemistry using diaminobenzidine as chromogen followed by counterstaining with Neutral Red as described before (Sooksawate et al., 2013). In brief, the slices were dipped into 0.3% Triton-X in phosphate buffer solution (PBST) containing 5% skimmed milk at room temperature and then with a rabbit anti-GFP antibody (1:2,000; Life Technologies, Japan) in PBST for 16 h at 4 °C. The sections were washed in PBST and incubated in biotinylated goat anti-rabbit IgG (1:200; Vector Laboratories, USA) and then ABC-Elite (1:200; Vector laboratories, USA) and visualized with diaminobenzidine (1:10,000; Wako, Japan) containing 1% Nickel sodium ammonium and 0.0003% H2O2 in Tris-buffered saline. Photomicrographs of the fluorescence were taken using fluorescent microscope (Akioplan 2, Zeiss, Oberkochen, Germany), histological slices were taken using light microscope (BZ-9000, BZ-X710, Keyence, NJ, USA) and the drawing were performed using a light microscope (BX-51, Olympus, Tokyo, Japan).

### Statistical analysis

Data are expressed as mean ± standard error of the mean (SEM). Significance was tested by Student’s unpaired *t-*test (Two-sided), and a p value < 0.05 was considered to be significant.

## Supporting information

Supporting information includes five video and can be found with this article online at https://doi.org/

**Supplementary movie S1. Head turn response**.

Pathway selective optogenetic activation-induced head turn (category 4) of mouse in the crossed pathway group to blue laser (50 ms duration) in the closed box.

**Supplementary movie S2. Circling response in closed box**.

Pathway selective optogenetic activation-induced circling response (category 1, >180°) of mouse in contralateral (crossed) pathway group to blue laser (50 ms duration, 10 Hz, 20 trains) in the closed box.

**Supplementary movie S3. Circling response in open platform**.

Pathway selective optogenetic activation-induced circling response (category 1, >180°) of mouse in ipsilateral (uncrossed) pathway group to blue laser (50 ms duration, 10 Hz, 20 trains) on the open platform. This mouse was the same one as in supplementary movie S2.

**Supplementary movie S4. Flight response in closed box**.

Pathway selective optogenetic activation-induced flight response (category 6) of mouse in ipsilateral (uncrossed) pathway group to blue laser (50 ms duration, 10 Hz, 20 trains) in the closed box.

**Supplementary movie S5. Retreat response in open platform**.

Pathway selective optogenetic activation-induced retreat response (category 5) of mouse in ipsilateral (uncrossed) pathway group to blue laser (50 ms duration, 10 Hz, 20 trains) on the open platform. This mouse was the same one as in supplementary movie S4.

